# G protein-coupled receptor SmGPCR9 interacts with neuropeptides and controls spermatogenesis in *Schistosoma mansoni*

**DOI:** 10.64898/2026.03.19.712866

**Authors:** Saranya Geetha, Simone Haeberlein, Steffen Hahnel, Xuesong Li, Daniel Sprague, Yuri K. Peterson, Shafqat Shabir, Franco H. Falcone, Moritz Buenemann, Christoph G. Grevelding

## Abstract

Schistosomiasis is a neglected tropical disease caused by parasitic flatworms of the genus *Schistosoma*, impacting hundreds of millions of people and animals globally. Disease pathology primarily originates from host immune responses to parasite eggs, which are produced only when female schistosomes are continuously paired with males. Past research focused on pairing-dependent female sexual maturation, while scarce data exist for the malés reproductive biology. In this study, we characterized the G protein-coupled receptor Sm*gpcr9* (Smp_244240), an orphan Class A (Rhodopsin-like) GPCR with a testis-preferential and pairing-influenced expression profile in *S. mansoni* males. Previous bulk RNA-seq analyses of adult worms and their isolated gonads revealed that Sm*gpcr9* belongs to a subgroup of GPCR genes with abundant testis-preferential and pairing-influenced transcript levels in males but low and extremely low expression in unpaired and paired females, respectively. This male-/unpaired female-biased expression pattern mirrors that of neuropeptide (*npp*) genes of *S. mansoni* such as Sm*npp26* and Sm*npp41*.

In a deorphanization approach using yeast-two-hybrid analyses, GPCR internalization experiments, bioluminescence resonance energy transfer assays, and by modeling and docking analyses, we provide first evidence that both NPPs can interact with SmGPCR9. Furthermore, we optimized a GPCR RNAi approach and achieved efficient transcript knockdown (> 90%) enabling robust functional characterization of Sm*gpcr9*. Following RNAi, physiological and morphological analyses revealed that SmGPCR9 regulates key aspects of male reproductive biology like testis morphology and spermatogenesis. Remarkably, ovary structure and egg production were also affected in paired females post RNAi. We observed similar phenotypes plus motility constraints and reduced stem-cell proliferation in both sexes upon RNAi of Sm*npp26* and Sm*npp41.* In all cases, RNAi downstream analyses by RT-qPCR of marker genes substantiated the observed phenotypic effects.

These results strongly indicate the importance of SmGPCR9, SmNPP26, and SmNPP41 for spermatogenesis and further physiological processes in male and female *S. mansoni*.

**Author Summary:** Research of the reproductive biology of schistosomes focused mainly on females so far, which upon pairing sexually mature to produce eggs that are important for the life cycle maintenance but also for the pathogenesis of schistosomiasis, the infectious disease caused by these parasites. We investigated a yet unknown G protein-coupled receptor, Sm*gpcr9*, which showed a testis-preferential and pairing-influenced expression profile in *Schistosoma mansoni* males. To this end, we optimized an RNA interference (RNAi) approach for knockdown analysis, identified neuropeptides (NPPs) as potential ligands by different biochemical approaches and modeling and docking analyses, and we investigated the roles of SmGPCR9 and two interacting NPPs, SmNPP26 and SmNPP41, by physiological, microscopical, and molecular techniques. Our results strongly suggest that SmGPCR9 and both NPPs regulate spermatogenesis. Furthermore, we detected effects on ovary morphology, egg production, and stem-cell proliferation of paired females post RNAi. Taken together, we deorphanized SmGPCR9 and showed for the first time the essential role of a so far uncharacterized GPCR and two interacting neuropeptides for spermatogenesis. Our results shed first light on spermatogenesis regulatory processes controlled by GPCRs and neuropeptides in male *S. mansoni* and thus expand our understanding of the roles of GPCR-NPP signaling for schistosome reproductive biology.

## Introduction

In eukaryotes, GPCRs represent a large and assorted group of transmembrane receptors. Due to their diversity, GPCRs are involved in a great variety of physiological processes, including visual, olfactory and taste perception, neurotransmission, hormonal regulation, immune response, and reproduction, including spermatogenesis [1–9]. To fulfil their roles, GPCRs can interact with different classes of ligands like hormones, neurotransmitters, gases, volatile compounds, biogenic amines, and neuropeptides [10–12]. In human, Class A (Rhodopsin) is the most diverse subfamily of GPCRs. They comprise hormone, neuropeptide, neurotransmitter, and light receptors, which are ligand-activated to interact with guanine nucleotide-binding (G) proteins for signal transduction [13–19].

In parasite research, GPCRs have come into focus mainly due to their druggability and the necessity to find new treatment options for parasitic diseases [20–24]. Beyond that, only few studies have addressed the biological function of GPCRs in protozoan and metazoan parasites so far [20, 25–32]. Schistosomes are parasitic platyhelminths and responsible for schistosomiasis (bilharzia), a neglected tropical disease and zoonosis, which affects humans and animals mostly in the global south [33–34]. Despite its global burden, schistosomiasis is treated solely with the calcium channel antagonist praziquantel (PZQ), a drug effective mainly against adult worms but not juveniles or eggs. Clinical symptoms and pathology of schistosomiasis are induced by eggs, which are produced by paired females in the bloodstream of vertebrate hosts. In intestinal schistosomiasis, part of the eggs find their way into the liver causing chronic granulomatous inflammation and fibrosis as main symptoms [34–35]. Egg production in turn is only achieved upon completion of differentiation of the female gonads (ovary and vitellarium), which depends on constant pairing with a male partner [36–38]. By inducing mitoses and differentiation processes in the reproductive organs, males play an essential role in regulating female sexual maturation [37–41]. Pairing even controls the expression of female-specifically expressed genes with functions in the gonads [42–44]. Past studies excluded sperm as a component for female maturation because anorchid males, paired with females, induced maturation and egg production [45–46]. Hormonal factors or peptides delivered by males were also postulated [39, 45–49, 50–51]. Recently, a research team identified a male-derived peptide-based pheromone, β-alanyl-tryptamine, as a first important factor regulating female sexual maturation [52]. The latter is supported by host vitamins, which in cooperation with the male-influence contribute to gonad differentiation in the female during pairing [44].

Bulk RNA-seq analyses of paired (bisex male, bM) and unpaired (single-sex male, sM) *S. mansoni* males and females (paired, bisex female, bF; unpaired, single-sex female, sF) and their gonads provided much information about sex-dependently, pairing-influenced and/or gonad-specifically or -preferentially expressed genes, including GPCRs [42]. Among these, few GPCRs showed testis-preferential and pairing-influenced expression in males but low abundant transcripts in sF, and nearly no expression in bF [53]. Remarkably, the vast majority of *npp* genes appeared to be regulated in a sex- and pairing-dependent manner with a male/unpaired female bias [42, 54]. Among the *gpcr* genes exhibiting a gonad-preferential and pairing-influenced expression pattern in males was Smp_244240, which we renamed Sm*gpcr9*. This orphan receptor belongs to the Class A (Rhodopsin) subfamily of GPCRs in *S. mansoni* [42, 53]. Here, our team provides first evidence for the interaction of SmGPCR9 with specific neuropeptides, SmNPP26 and SmNPP41. Functional analyses of all three genes demonstrated (i) testis-preferential (Sm*gpcr9*) and neuronal (Sm*npp26*, Sm*npp41*) transcript occurrence, (ii) roles in testis morphology and spermatogenesis, (iii) ovary morphology, (iv) egg production, and (v) stem-cell proliferation.

## Results

### Sm*gpcr9* is testis-preferentially expressed and potentially interacts with neuropeptides

Based on previous RNAseq data, we identified Sm*gpcr9* (Smp_244240) as a testis-preferentially and pairing-dependently transcribed gene in male *S. mansoni* [42, 53–55]. This expression pattern was confirmed by RT-PCR and indirectly by single-cell RNA-seq data of *S. mansoni*, which independently showed testes-preferential transcription of Sm*gpcr9* but also abundant transcript occurrence in the neuronal cell-cluster 3 (NC3) of males [56] (S1 Fig). Furthermore, based on phylogenetic analysis Sm*gpcr9* was categorized as a member of the Class A [Peptide (NPY, F, FF-like)] GPCR receptor subfamily in *S. mansoni*, and it is located on the sex chromosomes [53, 55]. These findings led us to characterize Sm*gpcr9* in more detail such as identifying potential interaction partners.

With the help of the membrane-anchored ligand and receptor yeast two-hybrid system (MALAR-Y2H) established in our lab [57], we searched for neuropeptide ligands (NPPs) of *S. mansoni* as potential interaction partners of Sm*gpcr9*. To this end, we screened an in-house ligand library comprising 47 *S. mansoni* neuropeptides (SmNPPs). By robust yeast growth on SD/Trp⁻ Leu⁻ His⁻ Ade⁻ selection plates, Sm*gpcr9* showed strongest interaction with the neuropeptides SmNPP2a, SmNPP5b, SmNPP15b, SmNPP26a, SmNPP32.2, and SmNPP41. Weaker growth was observed with SmNPP1a and SmNPP24, while all other NPPs resulted only in moderate or no interactions [57] (S2 Fig). Hence, we selected the mentioned SmNPPs for agonist internalization experiments to obtain additional hints for interaction and functional responses.

### Codon-optimized Sm*gpcr9* internalized in response to specific NPPs in HEK cells

For enhancing the expression in human cells, the Sm*gpcr9* coding sequence (CDS) was human codon-optimized (S3 Fig), tagged with dsRed and cloned into a modified pcDNA 3.1 vector backbone containing the mCitrine coding sequence as additional tag. With this construct, we transfected HEK293-6E suspension cells. To assess receptor-NPP interaction, soluble synthetic SmNPPs were added to the transfected cells. Internalization of the fluorescently labeled receptor upon ligand exposure served as a proxy for potential NPP binding and SmGPCR9 activation. Notably, except for SmNPPs 1a and 24, which already failed to induce growth and β-Gal activity in the former yeast experiments, all other tested NPPs induced internalization of SmGPCR9, as evidenced by the intracellular distribution of signals (Fig. 1 A-E; S4 Fig). These results suggested that the tagged version of SmGPCR9 is functionally expressed in HEK cells and bound by selected SmNPP ligands, which corresponded to the MALAR-Y2H data.

**Figure 1.**
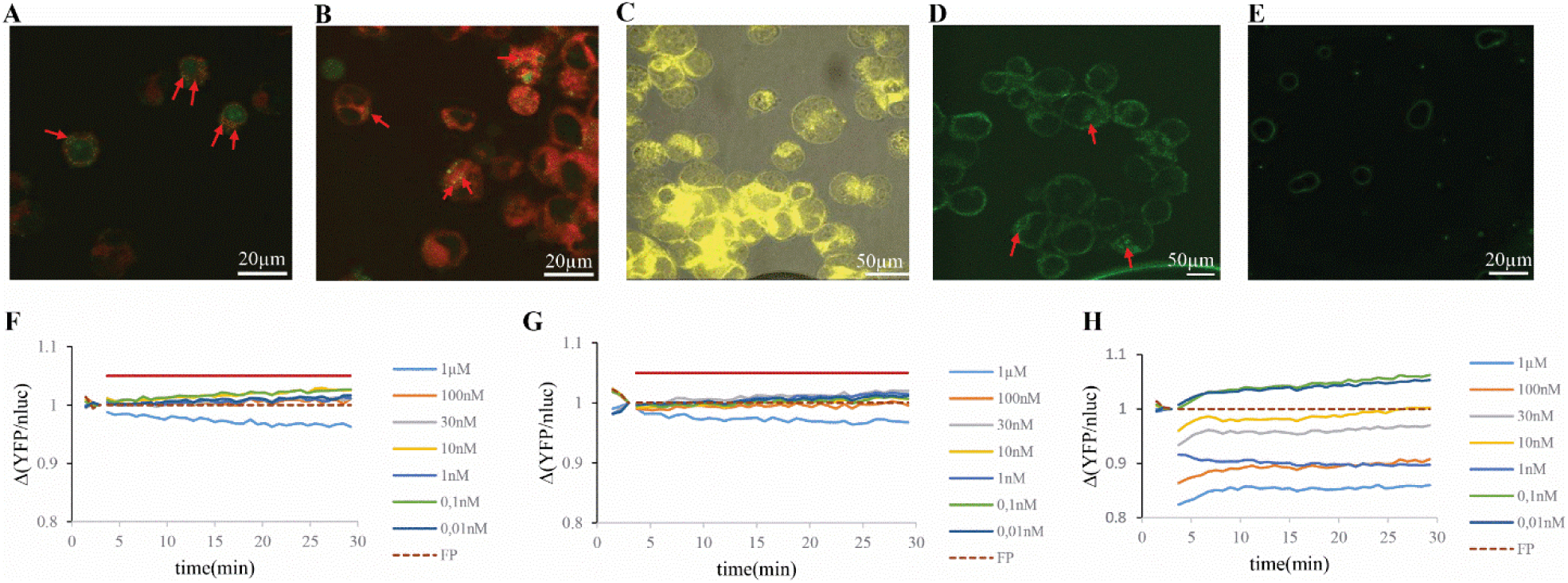
Cell internalization experiments and BRET assays identify SmNPP26 and SmNPP41 as SmGPCR binding partners. **A-B**, Selected examples of agonist-induced internalization experiments of the labelled version of SmGPCR9 (pcDNA_mCitrine_GPCR9_dsRed [N-terminal]) in HEK 293-6E cells upon stimulation with soluble SmNPP26b (A) and SmNPP41 (B), respectively (see also S4 Fig). In merged images, labelled SmGPCR9 appears as orange/yellow puncta (mCitrine + dsRed channels), indicated by red arrows. The occurrence of internal signals indicated ligand-induced internalization of the activated receptor. **C**, Negative control showing an overlay image of the labelled version of SmGPCR9 cells without ligand; as expected, signals are mainly found in the membrane. **D**, Positive control showing the internalization of the M2 muscarinic acetylcholine receptor following carbachol stimulation (green signals in the cytoplasm indicated by red arrows). **E**, Negative control for D showing M2-expressing HEK293 cells without carbachol stimulation. **F-G**, Results of Gαq recruitment assay showing positive responses for SmNPP26b (F) and SmNPP41 (G), respectively. The dashed line indicates the FRET buffer control used for normalization in this assay. **H**, Positive control showing the muscarinic acetylcholine receptor activated by carbachol. Scale bars as indicated. Experiments were performed in two independent biological replicates.

### Gαq-signaling assay indicated SmGPCR9 activation by SmNPP26 and SmNPP41

Next, we performed BRET (Bioluminescence Resonance Energy Transfer) assays between Gαq-nluc and Gβγ-YFP to find further evidence for SmGPCR9 interaction with the selected NPPs. Out of the candidate NPP panel, only SmNPP26 and SmNPP41 triggered reproducible BRET signals, which indicated Gαq activation by SmGPCR9. In both cases, we observed a decrease in BRET signal intensity at a concentration of 1 µM of the appropriate SmNPP (Fig. 1 F-H; S5 Fig A). To investigate whether the presence of the tagged receptor version may have influenced receptor function, we generated an additional construct encoding the untagged version of SmGPCR9 and cloned this variant directly into pcDNA3.1. HEK293T cells transfected with this construct were subjected to the same BRET assay protocol (S5 Fig B). However, we observed no significant differences of these results to the previous ones. Again, SmNPP26 and SmNPP41 exhibited activity at 1 µM, thus confirming the results with the tagged receptor version.

### Potential binding poses of SmNPP26 and SmNPP41 in GPCR9

Since we obtained accumulating evidence for SmGPCR9 interaction with SmNPP26b and SmNPP41, we additionally employed induced-fit molecular docking to predict binding poses of both SmNPPs in the extracellular domain of SmGPCR9. To this end, we first generated a homology model of SmGPCR9, based on the crystal structure of the human Neuropeptide FF receptor 2, PDB: 9M54. This template gave 74% coverage and 23% sequence identity to SmGPCR9 and resulted in the putative model (Fig. 2 A). Next, we prepared the two most likely NPP binding partners for docking. SmNPP26b and SmNPP41 did not allow homology modeling because of their small sizes. We attempted generating consensus conformers of SmNPP26b and SmNPP41 using a conformational search within the MOE computational suite. SmNPP41 returned 9 conformers that were practically identical, while SmNPP26b generated only a single conformer. Therefore, a single optimized conformer was used as ligand for flexible docking into the receptor.

**Figure 2.**
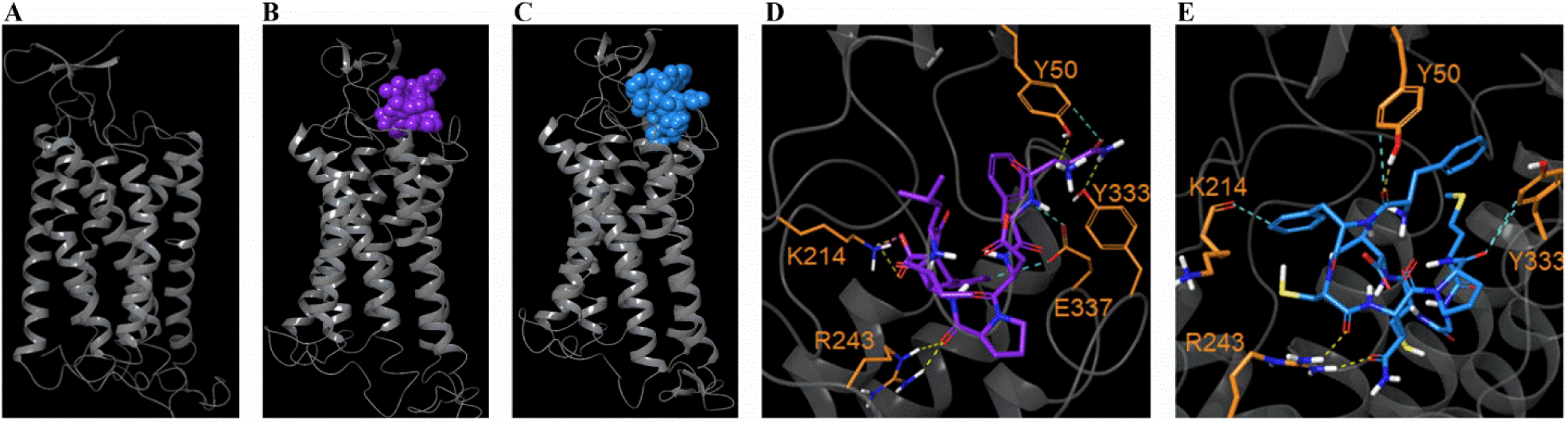
Molecular modelling and docking map binding areas of SmNPP26 and SmNPP41 in SmGPCR9. **A**, Molecular modeling of SmGPCR9. **B-C,** Space-filling model of SmNPP26b (purple), and SmNPP41 (blue) respectively in SmGPCR9 (grey) using induced-fit modeling. **D,** Predicted binding pose of SmNPP26b (purple) in GPCR9 (grey). Key interacting residues are labeled in orange. Clustering of the results showed a stable binding pose, in which SmNPP26 (purple) fits within an extracellular binding pocket of the receptor. SmNPP26 is predicted to be held into place by key hydrogen bonds to Y50, K214, R243, and Y333. In addition, it is predicted to form a salt bridge with K214, and aromatic hydrogen bonds with E337. Hydrogen bonds = yellow; aromatic hydrogen bonds = blue; salt bridge = purple. **E,** Predicted binding pose of SmNPP41 (blue) in GPCR9 (grey). SmNPP41 is also predicted to be held in place with hydrogen bonds to the side chains of Y50, R243, Y333 and an aromatic hydrogen bond to the backbone carbonyl of K214 instead of E337 in SmNPP26.

Having generated a receptor model and a stable conformer of the putative ligands, we then employed induced-fit docking, an approach where both the ligands and receptor are flexible, to determine possible binding poses. After clustering of the results, we obtained stable binding poses for both SmNPP26b (Fig. 2 B) and SmNPP41 (Fig. 2 C) where both SmNPPs (purple and blue, respectively) tightly fit within an extracellular binding pocket of SmGPCR9 (grey). Predicted binding free energies were favourable for both peptides, with SmNPP26b exhibiting a docking score of −7.38 kcal/mol and SmNPP41 showing slightly stronger predicted binding at −8.02 kcal/mol. The extracellular binding domain is structurally constrained and well defined, consistent with canonical GPCR architecture and the homology template. GPCRs typically bind ligands near the centre of the transmembrane helical bundle, with an extracellular domain composed of beta-sheet structures from the amino terminus and the E4 loop (between TM4 and TM5). This architecture occludes a large portion of the extracellular surface, thereby creating a discrete and well-defined ligand-binding pocket that accommodates both SmNPP peptides. The peptide is predicted to be held into place *via* similar key hydrogen bonds in both cases. For SmNPP26, the neuropeptide is predicted to form hydrogen bonds with Y50, K214, R243, and Y333 (Fig. 2 D). SmNPP26 was also predicted to form a salt bridge with K214, and aromatic hydrogen bonds with E337. SmNPP41 was also predicted to be held in place with hydrogen bonds to the side chains of Y50, R243, Y333 and an aromatic hydrogen bond to the backbone carbonyl of K214 (Fig. 2 E). These interactions, combined with favourable hydrophobic and hydrophilic environments, seem to be key for productive binding of both neuropeptides to SmGPCR9. Superposition of the top-scoring docked poses for SmNPP26b and SmNPP41 revealed a root-mean-square deviation (RMSD) of 2.85 Å, indicating highly similar binding modes. Notably, the proline and two phenylalanine residues in both peptides adopt closely overlapping positions within the binding pocket, suggesting conserved structural determinants for receptor engagement. Importantly, these similarities emerged from independently optimized best-scoring poses for each peptide, rather than from constrained alignment, further supporting a shared and robust binding mode.

### Sm*gpcr9* transcripts localized in testes and Sm*npp26/*Sm*npp41* transcripts in neuronal cells

For further characterization, we performed whole-mount *in situ* hybridization (WISH) to localize the transcripts of Sm*gpcr9*, Sm*npp26*, and Sm*npp41* in pairing-experienced (bisex) males (bM), pairing-experienced (bisex) females (bF), pairing-inexperienced (single-sex) females (sF), and pairing-inexperienced (single-sex) males (sM). Sm*gpcr9* transcripts were mainly found in testes and weakly throughout the body in a stripe-like, punctiform pattern (Fig 3 A-C). The latter may result from Sm*gpcr9* expression in neuronal cell cluster 3 (Ncc3), as indicated by single-cell RNA-seq analysis [56] (S1 Fig). The WISH signals confirmed in more detail our previous results for Sm*gpcr9* transcript localization using a classical *in situ* hybridization technique of sections of male worms [58, 59], which at that time failed to detect neuronal expression.

**Figure 3.**
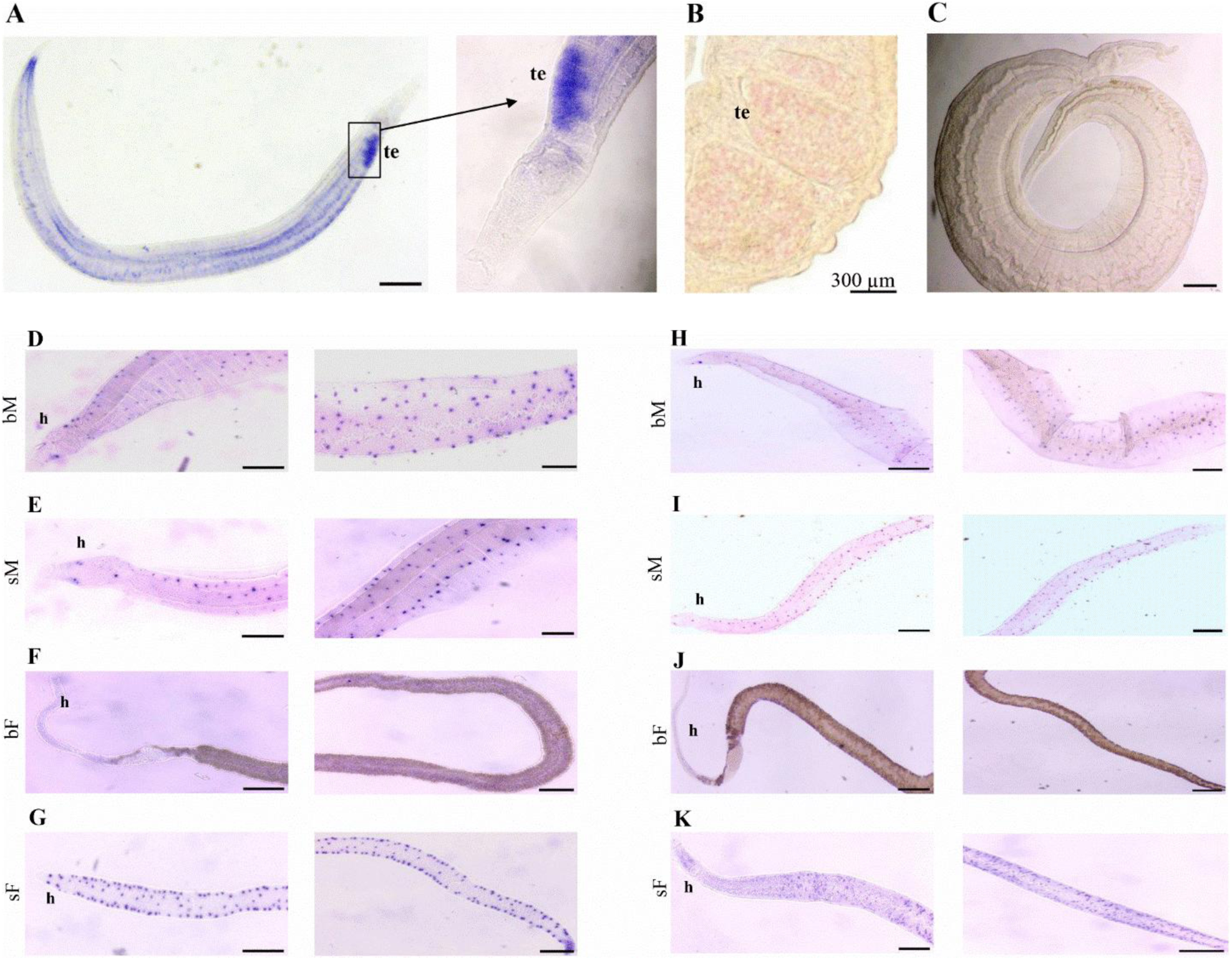
WISH localized Sm*gpcr9* transcripts mainly in testes, and Sm*npp26/*Sm*npp41* transcripts in neuronal cells. **A**, WISH result showing transcript localization of Sm*gpcr9* in male *S. mansoni* with magnification of the testes showing its typical lobular structure and strong signals; the right part is a close-up of the testes area shown in the left part. **B**, Sm*gpcr9* transcript localization (red signals) in the testes using a classical *in situ* hybridization technique of sections of male worms (technical details in 58). **C**, sense probe as negative control showed no signals. **D-G**, WISH result of Sm*npp26* transcripts in neuronal cells in the head parts and along the bodies of paired (bisex) males (bM; **D**); **E**, unpaired (single sex) males (sM); **F**, paired (bisex) females (bF; nearly no signals); **G**, unpaired (single sex) females (sF). **H-K**, Localisation of Sm*npp41* transcripts in neuronal cells of paired (bisex) males (bM; H); **I**, unpaired (single sex) males (sM); **J**, paired (bisex) females (bF; nearly no signals); **K**, unpaired (single sex) females (sF). If not indicated by other sizes, scale bars: 200 µm. Abbreviations: te, testes; h, head (anterior) part. Images are representative of 3-5 worms per group.

For the neuropeptides Sm*npp26* and Sm*npp41*, hybridization signals in males (bM and sM) exhibited punctiform distribution throughout the worm bodies (Fig 3, D-E and H-I), which is typical for neuronal expression patterns [32, 60, 61]. Cell atlas data demonstrated a wider transcript occurrence of Sm*npp26* and Sm*npp41* in many male tissues but dominant transcript occurrence in neuronal clusters 4 (Sm*npp26*) and 8 (Sm*npp41*), respectively [56] (S6 Fig). In females, distinct expression differences were observed between sF and bF groups. In sF, WISH signals of both *npp* transcripts prominently occurred in punctiform patterns, similar to their occurrence in males but more concentrated at the body edges (Fig 3, F-G and J-K). In contrast, signals were markedly reduced or absent in bF. These findings perfectly correspond to the previous bulk RNA-seq data of male and female *S. mansoni*, which showed abundant transcripts of Sm*npp26* and Sm*npp41* in adult worms but not in their gonads (S6 Fig). Furthermore, transcript levels in females were found to be pairing-dependent, with clearly higher mRNA levels of both *npps* in sF compared to bF. We confirmed these transcription patterns by RT-qPCR (S7 Fig). Sense probes used as negative controls for each target gene yielded no detectable signal, confirming probe specificity. Furthermore, control probes for known marker genes validated the integrity and accuracy of the WISH procedure: Sm*tsp-2* transcripts were localized near the tegument, consistent with its known role as surface-associated tetraspanin [62], while Sm*myst4* transcripts were restricted to the vitellarium, as expected for a female reproductive tissue marker [32] (S8 Fig A-B).

### RNAi against Sm*gpcr9*, Sm*npp26*, and Sm*npp41* influenced morphology and physiology of adult *S. mansoni*

In contrast to effectively knocking-down *npp* genes in *S. mansoni* by RNAi, the Sm*gpcr9* transcript level was reduced only about 50% with the standard method used in our lab (using a single dsRNA sequence per target gene) [44, 60, 63] (S9 Fig A). This comparatively low RNAi efficiency resulted in hardly reproducible testes phenotypes of treated worms *in vitro* (not shown). To overcome this limitation, we designed a two-probes/per-target-*gpcr* (tp/pt-*gpcr*) RNAi approach choosing two different sequence parts of Sm*gpcr9* for subcloning and dsRNA synthesis. SiRNA-finder (si-Fi) analysis supported the selection of suitable sequences, an approach which had been successfully applied before to optimize target-sequence selection by predicting potential off-target genes and to reduce or exclude off-target effects in *S. mansoni* [63, 64]. For tp/pt-*gpcr* RNAi, we used a concentration of 15 µg/mL per dsRNA, i.e. 30 µg/mL total amount of dsRNA for Sm*gpcr9*, which resulted in an increased knockdown (KD) efficiency of 90% ± 3% (S9 Fig B). For RNAi against Sm*npp26*, and Sm*npp41,* we treated worms with 15 µg/mL (7.5 µg/mL each of the two dsRNAs per gene) resulting in KD efficiency of > 90% in each case (S9 Fig C, D). To investigate RNAi effects for these three genes, we treated adult *S. mansoni* with the appropriate dsRNAs *in vitro* over a 15-day period.

By bright-field microscopy, we observed various phenotypic changes after dsRNA-treatment, especially in both *npp* RNAi groups. These phenotypes comprised curled and shrunken worm bodies, diminished motility, and the production of morphologically aberrant eggs. While RNAi of Sm*gpcr9* caused no obvious morphological changes, Sm*npp26* RNAi worms exhibited curling and shrinkage of the worm body. A similar but weaker phenotype was seen after Sm*npp41* RNAi (Fig 4).

**Figure 4.**
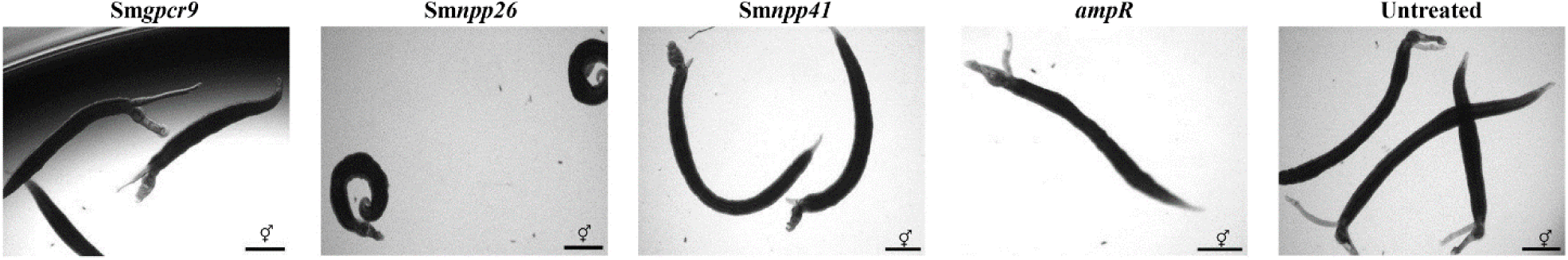
RNAi against all three target genes revealed morphologic changes of *S. mansoni* couples. Bright-field microscopy of *S. mansoni* couples revealed morphologic changes after treatment with dsRNAs targeting Sm*gpcr9*, Sm*npp26*, and Sm*npp41*, as indicated. In contrast to the controls (untreated and *ampR* dsRNA-treated), worm couples of the treatment groups (Sm*npp26*, and Sm*npp41*) were curved and/or curled, and they showed a shrinkage in size. Except slightly reduced motility after 12 d of dsRNA treatment (see text and Supplementary figure S10), no further obvious phenotypic changes were observed following Sm*gpcr9* RNAi. Experiments were performed in triplicates (n = 3). Scale bars = 100 µm.

For all three genes, RNAi had no influence on pairing stability within the observation period. In all three cases, however, worm motility diminished along the treatment period. Here, Sm*npp26* and Sm*npp41* RNAi showed significantly stronger effects already at day 3, whereas Sm*gpcr9* RNAi caused decreasing motility not before day 12 (S1 movie, S10 Fig). Although the total number of eggs was similar among all treatment groups, egg morphology differed. In all target gene groups, the number of abnormal eggs significantly increased at the end of the observation period (Fig 5).

**Figure 5.**
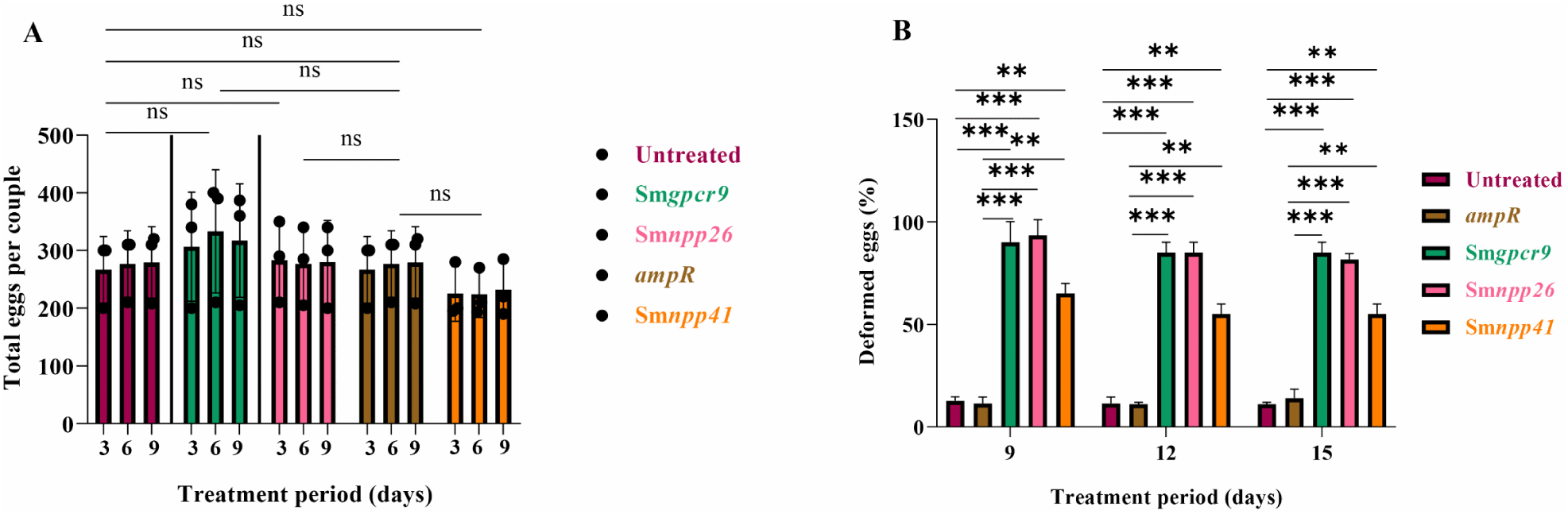
RNAi against all three target genes caused the production of deformed eggs. During the 15 days period of the RNAi experiment, eggs were produced *in vitro* in each dsRNA treatment group and in control worms (untreated, no dsRNA; *ampR*, irrelevant dsRNA, as indicated). **A**, Egg production (total amount) was similar in each dsRNA treatment group and in control worms. Compared to the other groups, a slight reduction was observed for worms of the Sm*npp41* RNAi group. **B**, Compared to the controls, the number of deformed eggs significantly increased in all RNAi groups from day 9 of treatment on. Significant differences were determined by t-test and indicated as: ***P < 0.001, **P < 0.01, *P < 0.05; ns, no significance. Experiments were performed in triplicates (n = 3).

In contrast to the controls, *in vitro*-laid worm eggs of the other treatment groups exhibited various defects including size reduction, missing spines, and/or no zygotes from day 9 forward (S11 Fig). These results reveal that RNAi against the three genes in focus caused effects on egg morphology over time but not on the total amount of eggs produced. These effects were independent of pairing stability, which was not influenced by RNAi, although worm motility was affected.

Following RNAi against all three genes, we investigated morphological changes in males and females by confocal laser scanning microscopy (CLSM). In males, *Smgpcr9* RNAi resulted in shrunken testicular lobes and a clear reduction in sperm content. In 3/5 males, sperm were completely absent whereas in the other 2 males, sperm content was very low. Similar testes phenotypes were seen after RNAi against Sm*npp26* and Sm*npp41*. In both control groups, untreated or treated with irrelevant dsRNA, none of these phenotypes occurred, and sperm vesicles were filled with differentiated sperm, as expected [65–66] (Fig 6 A).

**Figure 6.**
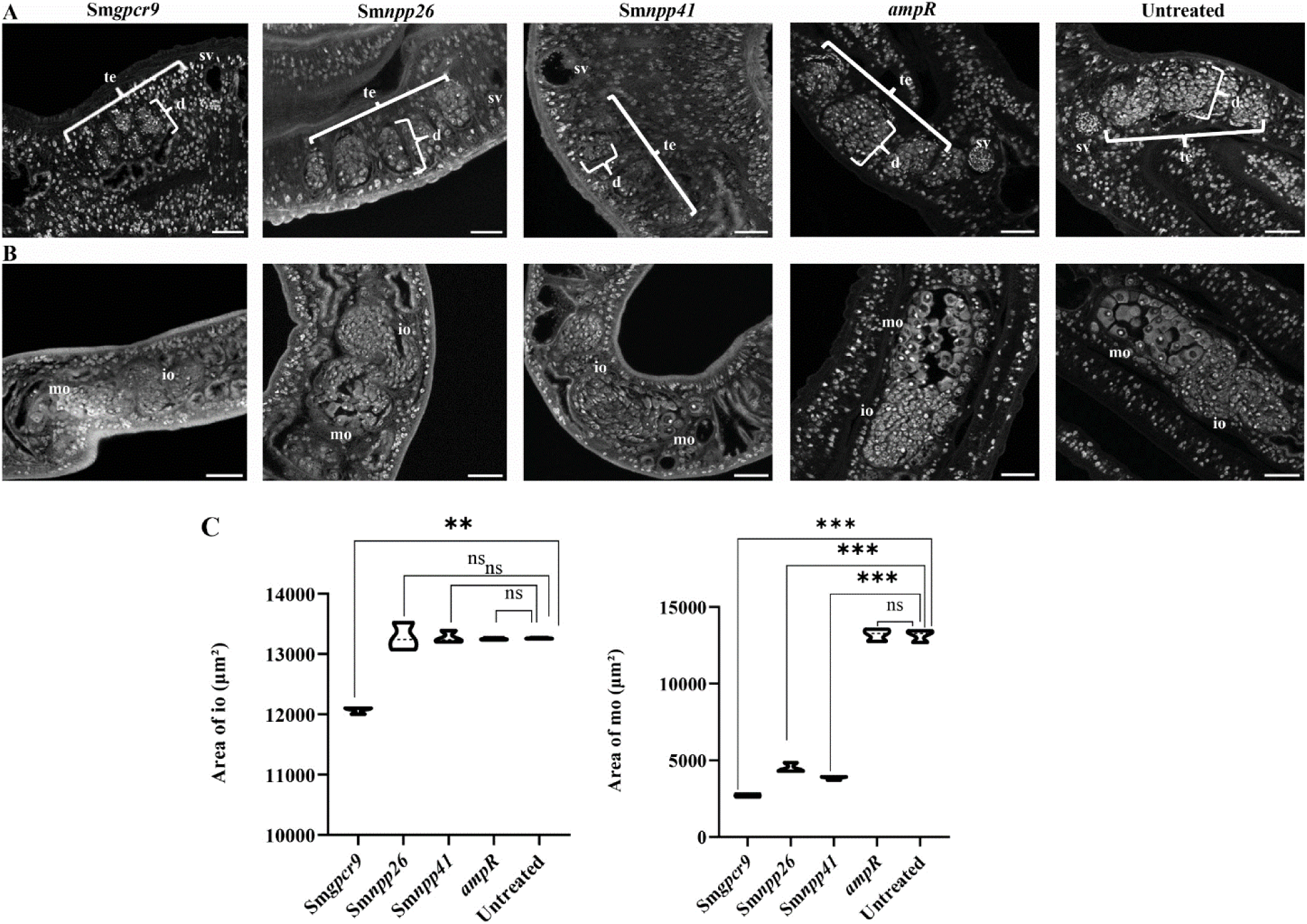
RNAi against all three target genes showed gonadal phenotypes in both genders. **A**, Following RNAi of all three target genes (as indicated), CLSM analyses of males revealed testes phenotypes such as shrunken testicular lobes with lower numbers of spermatogonia compared to both control groups (untreated, no dsRNA; *ampR*, irrelevant dsRNA). Also sperm production was diminished in all three target RNAi groups. None of these phenotypes occurred in the controls, in which sperm vesicles were filled with differentiated sperm, as expected (65-66). Scale bars = 50 µm. Abbreviations: te, testes; sv, sperm vesicle; d, diameter. **B**, In females, all three target RNAi groups showed reduced sizes of the ovaries, for Sm*gpcr9* with respect to both parts of the ovary, the posterior part containing mature oocytes and the anterior part containing immature, stem cell-like oogonia. Sm*npp26* and Sm*npp41* RNAi mainly diminished the anterior part of the ovary part and caused a reduction of mature oocytes. **C**, For all experimental groups (as indicated), we performed Image J-based determination of the sizes of the anterior part of the ovary containing immature, stem cell-like oogonia (Area of io; left diagram) and its posterior part containing mature oocytes (Area of mo; right diagram). **B-C**, As expected, both control groups exhibited large ovaries containing many mature oocytes (65-66). Significant differences were determined by t-test and indicated as: ***P < 0.001, **P < 0.01, *P < 0.05, ns, no significance; n = 5; scale bars = 50 µm. Abbreviations: mo, mature oocytes; io, immature oocytes. d, diameter; te, testes; sv, seminal vesicle; io, part of the ovary containing immature oocytes; mo, part of the ovary containing mature oocytes.

In paired females, Sm*gpcr9* RNAi resulted in smaller ovaries including reduced sizes of both the posterior part containing mature oocytes and the anterior part containing immature oogonia, which represent germinal stem cells (GSCs). Upon Sm*npp26* and Sm*npp41* RNAi, especially the size of the posterior part of the ovary was reduced as well as the number of mature oocytes. As expected, in females of both control groups, ovary sizes were normal, and the numbers of immature and matures oocytes were high [65–66] (Fig 6 B, C).

Next, we performed 5-ethynyl 2’-deoxyuridine (EdU) incorporation assays to find out whether the described RNAi phenotypes in both sexes are associated with disturbed stem-cell proliferation. In males, Sm*gpcr9* RNAi caused no significant alterations in the number of EdU-positive cells, neither in GSCs within the testes nor the somatic stem cells, the neoblasts, which occur along the worm body [67]. Thus, stem cell proliferation following RNAi appeared normal and was similar to that of the male control groups (treated with irrelevant RNA and untreated). In contrast, we observed a significant reduction in the number of EdU-positive GSCs and somatic stem cells for Sm*npp41* but not for Sm*npp26* following RNAi (Fig 7 A, S12 Fig A-C).

**Figure 7.**
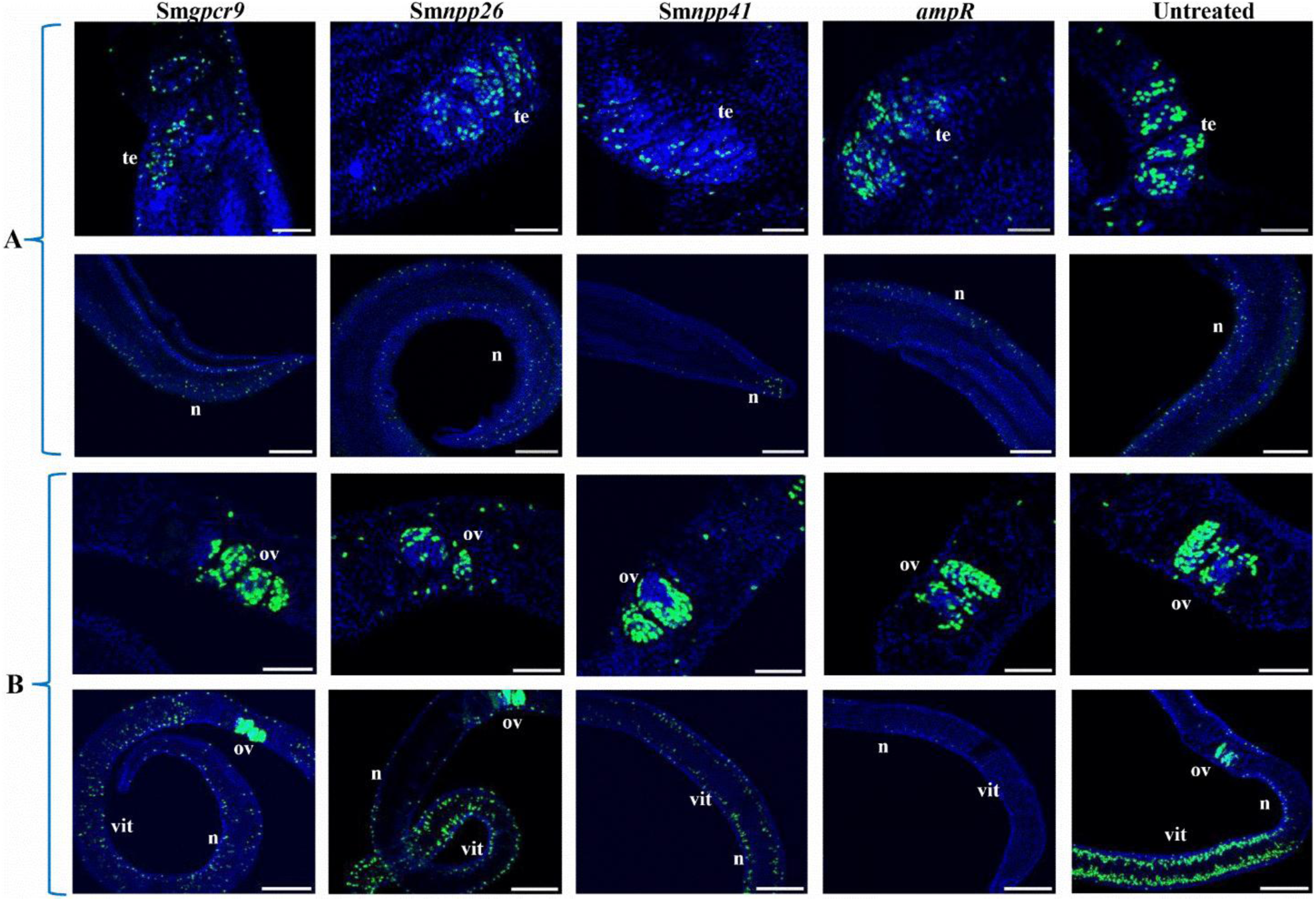
Gonadal stem-cell proliferation was affected upon RNAi against Sm*npp41* and Sm*npp26,* not Sm*gpcr9*. Results of EdU staining (green) of male (**A**) and female (**B**) *S. mansoni* after RNAi against the three target genes and the controls, as indicated. (**A**), In testes (te), a reduction of the number of EdU-positive gonadal stem cells (upper row) was found mainly for Sm*npp41* RNAi, a tendency of reduction for Sm*npp26* RNAi, but no reduction for Sm*gpcr9* and the control groups (untreated, no dsRNA; *ampR*, irrelevant dsRNA; see also S 12 Fig). Comparable reductions were observed for somatic stem cells (neoblasts (78), lower row), which are embedded in the parenchyma along the worm body. (**B)**, In females, a reduction of the number of EdU-positive gonadal stem cells (upper row) was found for Sm*npp26* and Sm*npp41*, but no reduction for Sm*gpcr9* and the control groups (see also S12 Fig). In the vitellarium, which belongs to the female gonad system (37), reduction of EdU-positive cells was mainly observed for Sm*npp41* (lower row). Positive vitelline stem cells are difficult to distinguish from neoblasts in the female. Unexpectedly, we found no staining in the *ampR* control. Hoechst was used as counterstain (blue). Experiments were performed with n = 3. Scale bars = 50 µm. Abbreviations: ov, ovary; vit, vitellarium; n, neoblast (somatic stem cell).

In females, reduced numbers of EdU-positive GSCs in the ovary were found for Sm*npp26* and Sm*npp41*, but not for Sm*gpcr9* and the control groups. Part of the female gonad is the vitellarium [37], for which we observed a clear reduction of EdU-positive cells mainly after Sm*npp41* RNAi. In contrast to the control without dsRNA (untreated), which expectedly showed much staining in the female gonads (ovary and vitellarium), we repeatedly (n = 3, five worms/n) found only few positive cells in the vitellarium following *ampR* dsRNA treatment. This finding was unexpected and suggests a yet undetected [63] vitellarium-specific effect of this “irrelevant” dsRNA on stem-cell activity in this organ (Fig 7 B, S12 Fig D).

In summary, CLSM and EdU-assay data indicated that Sm*gpcr9* RNAi caused smaller testes and reduced sperm production in males but without affecting stem-cell proliferation. In females, Sm*gpcr9* RNAi caused a similar gonadal phenotype, smaller ovaries without affecting stem-cell proliferation. Upon Sm*npp26 and* Sm*npp41* RNAi, similar testes phenotypes were observed in males with respect to reduced testes size and sperm production. Here, significant effects on stem-cell proliferation were found for Sm*npp41* but not for Sm*npp26*. In females, the size of the posterior part of the ovary and the number of mature oocytes were reduced in both the Sm*npp26* and Sm*npp41* RNAi groups, which was paralleled by significantly reduced stem cell proliferation in each case.

### RNAi against Sm*gpcr9*, Sm*npp26*, and Sm*npp41* affected the expression of selected genes

To solidify the observed phenotypic effects in both sexes and to find further hints for their molecular basis, we investigated the expression profiles of selected genes by RT-qPCRs. As starting material, we used couples treated for 15 days *in vitro* with the appropriate dsRNAs. At the end of the experimental period, couples were manually disconnected, and RNA was separately isolated from females and males. As qPCR control, Sm*letm1* was used as the reference gene based on its proven use for *in vitro* studies [68]. Following RNAi against each respective target gene, the expression levels of Sm*gpcr9* and Sm*npp26* were significantly reduced in both sexes. Whereas the expression of Sm*npp41* was also significantly reduced in males, it unexpectedly increased in females (Fig 8 A). With regard to the various RNAi phenotypes detected for Sm*gpcr9*, Sm*npp26,* and Sm*npp41*, we selected the following genes for downstream analyses. Concerning the suspected deficiency of sperm differentiation in males, we focused on tektins because they represent marker genes for sperm differentiation coding for structural components of sperm flagella, which are responsible for sperm motility [69–71]. Based on their expression profiles in RNAseq studies, we selected two annotated tektins of *S. mansoni*, Sm*tektin-a1* (Smp_343970) and Sm*tektin-2* (Smp_046410; https://parasite.wormbase.org) [55]. Both tektins showed similar transcript profiles as Sm*gpcr9* with a clear testis-bias of expression according to bulk RNAseq data [42] (S13 Fig A, B). Furthermore, scRNAseq data indicated the dominant occurrence of transcripts of both tektins in male testes but also in some Nccs and flame cells [56] (S13 Fig C, D).

**Figure 8.**
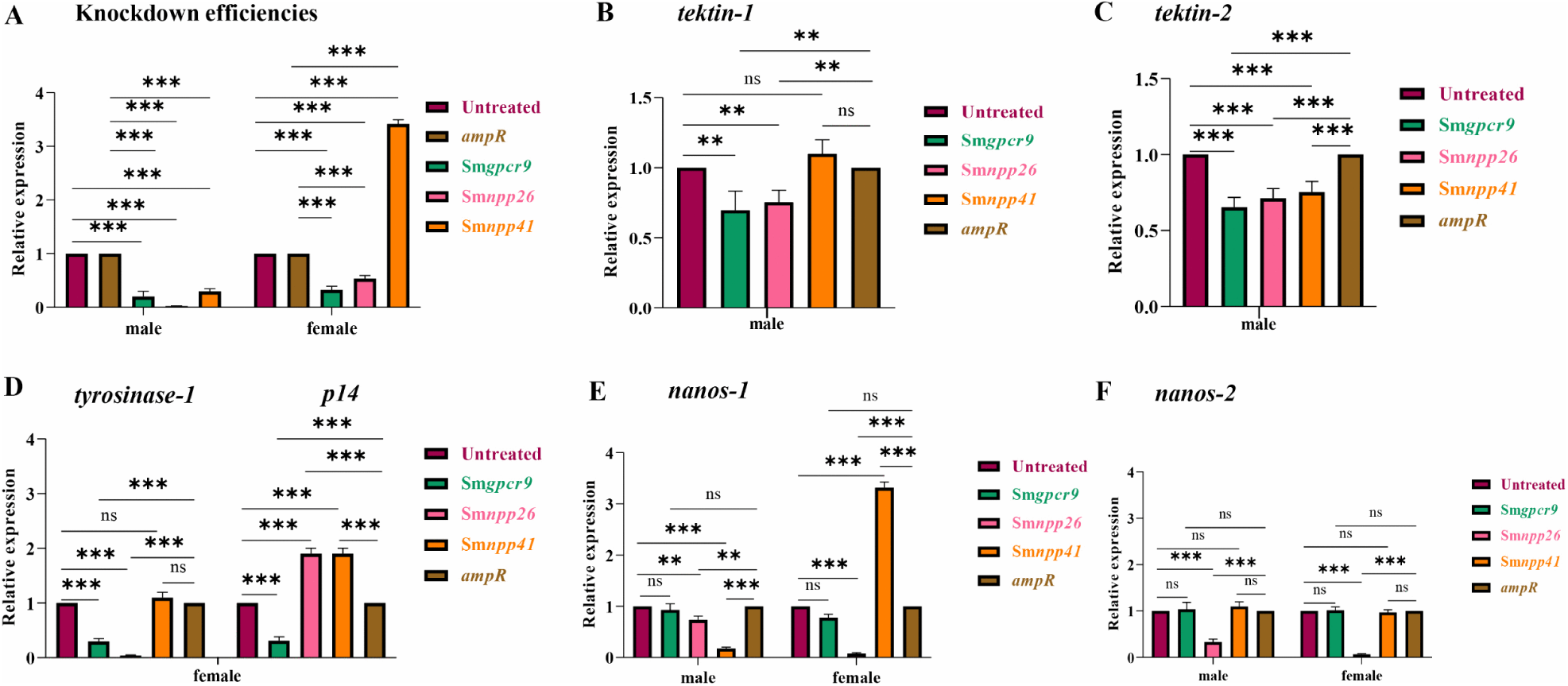
Marker gene expression was influenced following RNAi against the three target genes. **A**, Following RNAi against each respective target gene (as indicated), the transcript levels of Sm*gpcr9* and Sm*npp26* were significantly reduced in male and female *S. mansoni.* Similarly, the expression of Sm*npp41* was significantly reduced in males, however, it increased in females. Worms not treated with dsRNA or with irrelevant dsRNA (*ampR*) served as references. **B-C**, In male *S. mansoni*, significant reduction of the expression level of *tektin a1* was observed in the RNAi groups Sm*gpcr9* and Sm*npp26*, while for *tektin 2*, significant reduction of its expression level was observed in all RNAi groups (as indicated). **D**, In female *S. mansoni*, expression of both egg synthesis-related genes, *tyrosinase 1* and *Smp14*, was significantly downregulated following Sm*gpcr9* RNAi and Sm*npp26* RNAi. Sm*npp41* RNAi caused no significant change of the transcript level of *tyrosinase 1* but an increase of *Smp14* expression. **E**, The expression of *nanos-1* was significantly reduced in the Sm*npp26* RNAi groups of males and females, whereas Sm*gpcr9* RNAi caused no significant change of *nanos-1* expression. In the Sm*npp41* RNAi group, *nanos-1* expression was significantly reduced in males but strongly upregulated in females. **F**, For *nanos-2*, a significant reduction of expression was observed after Sm*npp26* RNAi in both sexes, whereas no difference occurred after Sm*gpcr9* and Sm*npp41* RNAi. Fold changes of gene expression levels between dsRNA-treated worms and untreated control worms were calculated using the 2-ΔΔCt method. Data are representative of the mean ± SEM of three separate experiments (n = 3). Significant differences were determined by t-test and indicated as: ***P < 0.001, **P < 0.01, *P < 0.05, ns, no significance.

Regarding diminished egg production, egg malformation, morphological changes in the female ovary, and the observed stem cell-proliferation effects, we analyzed marker genes that were shown before to be involved in egg production and stem-cell proliferation: (i) the egg-shell biosynthesis enzyme *tyrosinase 1* (Sm*tyr-1*; Smp_050270) [72], (ii) the egg-shell precursor gene Sm*p14* (Smp_131110) [73], (iii) the germline stem cell (GSC) marker gene *nanos-1* (Smp_055740) [56, 44], and (iv) the neoblast and GSC marker gene *nanos-2* (Smp_051920) [74].

Following Sm*gpcr9* RNAi, the transcript levels of both tektin genes were significantly reduced in males (Fig 8 B, C). Unexpectedly, we also observed significantly reduced transcript levels of Sm*tyr-1* and Sm*p14* in females (Fig 8 D). In contrast, expression levels of Sm*nanos-1* and Sm*nanos-2* were unaffected (Fig 8 E, F).

Sm*npp26* RNAi significantly reduced the expression levels of both tektin genes as well as *nanos-1* and *nanos-2* in males and females (Fig 8 B, C, E, F). Furthermore, we detected a significantly reduced expression level of Sm*tyr-1* but not Sm*p14* (Fig 8 D).

In males, Sm*npp41* RNAi significantly reduced the expression levels of Sm*tektin-2* and Sm*nanos-1* but not Sm*tektin-a1* and Sm*nanos-2* (Fig 8 B, C, E, F). In females, Sm*npp41* RNAi significantly enhanced the expression level of Sm*nanos-1* but not Sm*nanos-2*, Sm*tyr-1,* and Sm*p14* (Fig 8 D-F). With respect to the observed effects on marker gene expression in females, one must perceive the latter results with a certain degree of caution because the transcript level of Sm*npp41* appeared significantly upregulated in females after RNAi (Fig 8 A).

## Discussion

Usually, platyhelminths are hermaphroditic worm organisms that follow the principle of protandry (male gonad development precedes female gonad development) as one of two possible forms of sequential hermaphroditism [75–76]. Against this background, schistosomes are exceptional in having evolved sexual dimorphism during evolution. On their evolutionary path, however, absolute dioecy has not yet been achieved, because female schistosomes dependent on permanent pairing with their male partners to achieve complete sexual maturation, an obvious reminiscence to their protandric ancestors [77]. During constant pairing, the female lodges within the gynecophoral canal, a ventral groove formed by the male. This intimate contact can last for years as a prerequisite for mass production of eggs over time [36–41]. Since egg laying is a double-edged sword of schistosomes, ensuring life-cycle maintenance on the one side, and on the other causing pathology in the final host [34], female gonad development has been in the focus of many studies, while research did not pose enough importance on studying gonadal processes in males so far.

In the past, mainly single gene and signalling cascade analyses were performed to study schistosome male reproductive biology. In this context, a Fushi-tarazu factor-1 nuclear receptor was localized in the testis of male and ovary of female *S. mansoni* [78]. Mediated by excretory-secretory products, roles for extracellular signal-regulated kinase (ERK) and p38 mitogen-activated protein kinase (p38 MAPK) pathways were shown to affect cell proliferation, the tegument, the female gonads, and the testes of males [79]. Further, cytoplasmic protein tyrosine kinases of the Src (SmTK3), Src/Abl (SmTK6), Abl (SmAbl1, SmAbl 2), and Syk (SmTK4) families were shown to be expressed in male testes and to be involved in signal transduction pathways organizing the cytoskeleton in gonadal cells of *S. mansoni* [58, 80–82]. The *S. japonicum* ortholog of SmTK4 was shown to be important for gametogenesis also in this species [83]. Moreover, transcripts of the *S. japonicum* ortholog of pumilio, SjPum2, an RNA-binding protein, were found in testis and ovary. Sj*pum2* RNAi resulted in morphological alterations of both male and female gonads [84]. The latter phenotype was also demonstrated following RNAi of Sj*nanos1*, which is a gonadal stem-cell marker in *S. japonicum* [85]. Additional comprehensive analyses of bM versus sM by SAGE (Serial Analysis of Gene Expression) and microarrays unraveled additional and pairing-influenced roles for TGFβ-signaling, with Sm*fst* (follistatin) and Sm*bmp* (bone morphogenic protein) as genes expressed in testes and ovary [86]. Results of RNAseq analyses in *S. mansoni* finally suggested a complex scenario with even more molecular players that are differentially expressed upon pairing in both genders [42, 54]. A similar complexity was found in an RNAseq study of *S. japonicum* [29], which suggests the species-independence of these findings.

In *S. mansoni*, morphological studies by CLSM exhibited no substantial differences between testes of bM and sM, both have fully developed testicular lobes, spermatogonia, and fully differentiated sperm [65–66]. However, RNAseq identified 243 pairing-dependently expressed genes in the testis

[42], which suggests the existence of regulatory processes “underneath” the visually detectable morphological level [54]. Among these was Sm*gpcr9*, for which we confirmed transcriptional upregulation after pairing and dominant expression in the testes. Further WISH signals in the anterior “head” part and along the male body indicated additional expression in neurons. This coincided with scRNAseq data, which detected Sm*gpcr9* transcripts also in cells of the neuronal cluster 3 [56].

According to a previous classification based on phylogenetic analyses, Sm*gpcr9* represents a member of the Rhodopsin-like GPCR family, and it was connected to NPP signaling [53]. NPPs play important roles in reproduction as shown for different organisms like *S. mediterranea* [87–88], a free-living flatworm and closely related to schistosomes, and *Drosophila melanogaster* [89–91]. Therefore, we applied a deorphanization approach focusing on NPPs of *S. mansoni* that we had cloned in a MALAR-Y2H library. The initial library screening to identify potential interaction partners of SmGPCR9 identified five NPP candidates [57]. In this study, we restricted the number of putative interaction partners using GPCR internalization experiments, BRET assays, and modeling approaches of the two most likely candidates, SmNPP26b and SmNPP41. The latter approach included the first homology model of SmGPCR9 based on the human neuropeptide FF receptor 2, the most suitable template for this receptor. As expected, the homology model predicted seven transmembrane helices, which help to form an extracellular binding domain within which SmNPP26b and SmNPP41 can bind. Induced-fit docking predicted both NPPs to fit into this pocket, forming several favorable van der Waals interactions in addition to being held in place by key hydrogen bonds and salt bridges.

Besides Sm*gpcr9* transcripts in the testes, localisation experiments exhibited transcripts of all genes in various Nccs. Although the WISH signals for *npps* were mostly punctiform, the patterns differed in their positions. In males, Sm*gpcr9* transcripts occurred throughout the worm body along two parallel stripes, whereas the majority of Sm*npp26* transcripts occurred closer to the tegumental surface area. Sm*npp41* transcripts appeared more central, and patchy. These differences can be explained by scRNAseq results that detected Sm*gpcr9* transcripts (besides testes) mainly in Ncc3, Sm*npp26* transcripts dominantly in Ncc4, and Sm*npp41* transcripts dominantly in Ncc8. Comparing signal occurrence between sM and bM, it appeared as if the patterns were slightly different as well. Especially for Sm*npp41*, transcripts occurred closer to the edges of the worm body in sM, but they appeared more central in bM. This either suggests a change of SmNPP41 expression in different neuronal cells before or following pairing, or migratory capacities of involved neuronal cells. Indeed, dorsoventral and anterior-posterior migration has been described for vertebrates and invertebrates, and it can be part of differentiation processes [92–93]. We observed an even clearer pairing-dependent difference in females. Whereas Sm*npp26* and Sm*npp41* transcripts dominated in sF, occurring beneath the tegument (Sm*npp26*) or more central (Sm*npp41*), in both cases WISH signals were nearly absent in bF. These results confirm previous bulk RNAseq data of adult *S. mansoni* and their gonads [42] as well as RT-qPCR results (this study), which showed significantly reduced transcript levels of both *npps* in females after pairing. This finding supports the former hypothesis that reduced *npp* expression in bF might indicate a lower importance of female NPP activity after pairing. However, since NPP activities should be physiologically essential for both sexes, the question arises whether the male, after pairing, takes over “the neuronal power” over the female, and how? Part of this takeover is the reduced need for muscular activity of the female after she has lodged in the gynecophoral canal of the male, which after pairing acquires all locomotion activities of the couple. This belongs to the separation of labour arrangement of the schistosome couple, a hypothesized evolutionary advantage that distinguishes the exceptional, sexually dimorphic schistosomes from other hermaphroditic platyhelminths [94]. Another part of this takeover may deal with the reproductive biology of the female, which is governed by the male. Our RNAi-based functional analyses of all three genes provided evidence for this. For an easier overview of the results, we summarize and discuss the phenotypes for all genes in focus in the following sections.

### Smgpcr9

With the optimized tp/pt-*gpcr* RNAi approach, microscopy of dsRNA-treated worms showed no obvious morphological changes and no effect on pairing stability during the observation period of 15 d. However, motility decreased from d 12 on. In males, CLSM demonstrated shrunken testicular lobes and reduced sperm production upon Sm*gpcr9* RNAi, and EdU-assay results indicated normal stem-cell proliferation. These findings suggest that Sm*gpcr9* plays no role for GSC activities in the testes, but it does play a role for differentiation processes after stem-cell division that lead to spermatogonia. Support for this assumption was provided by RT-qPCRs demonstrating the downregulation of two tektins, Sm*tektin-a1* and *-2*, marker genes for sperm differentiation. In contrast, Sm*nanos-1/2*, marker genes for GSC proliferation were transcribed at the same level as the controls.

Unexpectedly, the number of deformed eggs significantly increased from d 9 on after Sm*gpcr9* RNAi, and CLSM of females exhibited smaller ovaries. Results of EdU assays, which showed no significant differences between the RNAi group and the controls, suggest that the gonadal phenotypes in females were independent of GSC proliferation. This corresponds to RT-qPCR results showing no effects on the expression levels of Sm*nanos-1/2*. In contrast, the transcript levels of Sm*tyr-1* and Sm*p14*, both differentiation markers for female vitellarium, as one part of the female gonad, were downregulated. This suggests that Sm*gpcr9* RNAi likely affected differentiation processes post stem-cell division also in bF. These effects were unexpected since Sm*gpcr9,* according to all RNAseq data for adult schistosomes available today, is a strictly controlled gene with a gonad-preferential, pairing-influenced, and Ncc3-associated expression profile in males. In females, bulk RNAseq data showed negligibly small transcript amounts, however, scRNAseq data showed transcript occurrence also in some further neuronal clusters (https://www.collinslab.org/schistocyte/search?gene=Smp_244240). The latter finding indicates a sex difference in Sm*gpcr9* transcription, which may contribute to the female phenotype. However, scRNAseq showed no Sm*gpcr9* transcripts in ovary or vitellarium. This could mean that the phenotype in females, which comprised higher production of deformed eggs, reduced ovary size, and decreased vitellarium marker gene expression, resulted either from an additional but indirect role of Sm*gpcr9* in males or from the expression of this gene in female Nccs. If the effect originates from males, then two scenarios are conceivable. Either sperm fluid transports factors have been produced downstream of SmGPCR9 activation in the testes and influence female gonad differentiation, or factors are transmitted to the female via excretory-secretory products (ESPs) during a pairing contact. Indeed, a recent study demonstrated a role of ESPs in male-female interaction and the reciprocal communication between the sexes [95].

### Smnpp26

Sm*npp26* RNAi resulted in strongly curled and constricted worm bodies from d 3 on, diminished motility, and induced the production of abnormal eggs although pairing stability was unaffected. In males, CLSM revealed smaller testicular lobes and the absence of differentiated sperm. EdU assays showed a trend but no significant reduction of stem-cell proliferation, while RT-qPCRs unraveled significantly reduced transcript levels of Sm*nanos-1/2*. Therefore, we cannot exclude a stem-cell effect in the male gonad. Nevertheless, the significantly reduced transcript levels of Sm*tektin-a1* and *-2* substantiate the CLSM phenotype and suggest diminished sperm differentiation, part of which could perhaps be explained by an additional RNAi effect on stem-cell proliferation.

CLSM of the female exhibited a smaller ovary, in which especially the posterior part was significantly smaller as was the numbers of mature oocytes. Although no obvious reduction of the part of the ovary containing immature oocytes was microscopically observed, EdU assay results indicated less GSC proliferation. This is paralleled by reduced transcript levels of Sm*nanos-1/2*. Since the transcript level of Sm*tyr-1* was also reduced, a differentiation effect on the vitellarium seems likely, which might explain the high number of deformed eggs because Sm*tyr-1* is responsible for egg-shell biosynthesis [72]. At the same time, Sm*p14* transcript levels were unchanged. This could be explained assuming that both genes are targets of parallel but different pathways controlling female sexual maturation upon pairing. In this theoretical scenario, Sm*tyr-1* would be a target of a Sm*npp26* pathway while Sm*p14* would be the target of another pathway.

### Smnpp41

Although not as intense as after Sm*npp26* KD, Sm*npp41* RNAi also caused early curling, body bending, and the production of abnormal eggs while maintaining pairing stability. In males, CLSM demonstrated similar testes phenotypes as for Sm*npp26* RNAi, however, the diameter of the testicular lobes appeared to be even smaller. EdU incorporation was significantly lower upon KD, which corresponds to the significant downregulation of Sm*nanos-1*, the GSC marker gene. Furthermore, the transcript level of Sm*tektin-a1* was significantly reduced, but not that of Sm*tektin-2*. Also in this case, we follow the idea that both genes might be targets of at least two different pathways, with Sm*tektin-a1* under the control of Sm*npp41*.

In females, the results are more difficult to interpret because Sm*npp41* transcript abundance was significantly higher following dsRNA treatment. Nonetheless, an ovary phenotype was obtained that resembled the one observed for Sm*npp26* RNAi female worms. EdU assays showed significantly reduced GSC proliferation, and marker gene expression analyses revealed either upregulation for Sm*nanos-1* or unchanged expression for Sm*nanos-2*. The Sm*tyr-1* transcript level was unchanged, and the level of Sm*p14* significantly upregulated. These findings indirectly support the theoretical scenario above that both genes could be targets of different pathways. Whether the upregulation of Sm*nanos-1* and/or Sm*p14* can explain the observed egg deformation in a gain-of-function/overexpression-like manner remains unclear at this stage of the analysis.

The observed upregulation of Sm*npp41* upon dsRNA treatment in females was unexpected, especially because the opposite (and expected downregulation) effect was found for males. RNAi has been discussed as an evolutionary conserved mechanism of prokaryotes defending against “hostile” RNAs sequences, such as mobile genetic elements or viral RNA [96]. In eukaryotes, evolution has adapted this innate defence system to coordinate complex gene regulation mechanisms. This can encompass epigenetic processes and chromatin with different outcomes, regulating transcription negatively and positively [97–99]. Against this background, the finding of opposing RNAi effects using the same dsRNA at the same time for paired male and female *S. mansoni* may be explained by regulatory mechanisms in the promoter region of the Sm*npp41* gene that differ between the sexes. As the level of Sm*npp41* transcripts is strongly reduced after pairing in females, opposite to males, it seems likely that epigenetic processes at the chromatin level might be involved. If this is the case, it is tempting to speculate that the used Sm*npp41* dsRNA may have different epigenetic partners in males and females to interact with at this promoter region. This could have caused the sex difference.

In summary, with SmGPCR9 we have identified the first GPCR of a parasitic flatworm with a proven role for spermatogenesis. Our molecular analyses suggest that SmGPCR9 controls sperm differentiation, probably at the level of primary or secondary spermatocytes. Upregulation of SmGPCR9 expression in males after pairing may reflect the need for reinforced sperm production. In addition, SmGPCR9 appears to be involved in processes regulating female sexual maturation, as part of the complex male-female interaction. This finding extends results of a previous study that demonstrated the importance of NPP-stimulated GPCR signaling in the neuroendocrine control of germ cell differentiation in planarians, free-living platyhelminths [87]. In fact, our deorphanization approaches identified two neuropeptides, SmNPP26 and SmNPP41, as potential interaction partners. Their functional characterization showed phenotypes in both sexes that overlapped with those found after Sm*gpcr9* RNAi in both sexes. This substantiates their role as potential SmGPCR9 ligands. Further phenotypes without intersection with the Sm*gpcr*9 KD results suggest that SmNPP26 and SmNPP41 probably bind also to other receptors. As a side note, the strong curling and motility phenotype of Sm*npp26* RNAi is remotely similar to that observed after PZQ treatment [100]. This indicates not only a further role of SmNPP26 in controlling neuromuscular activity, which might explain the curling phenotype, but also its role as candidate for target evaluation experiments with the aim to find urgently needed new drugs for schistosomiasis [101].

## Material and Methods

### Experimental design describing the objectives and design of the study as well as respecified components

To conduct the first functional characterization of Sm*gpcr9* (Smp_244240), an orphan Class A (Rhodopsin-like) GPCR of *S. mansoni*, we performed a deorphanization approach using data of a previous Y2H analysis (57), GPCR internalization experiments, BRET assays, and modeling and docking analyses. Furthermore, with the help of an optimized RNAi approach, we investigated the knockdown effects of Sm*gpcr9* and two NPPs in adult *S. mansoni in vitro*. This included morphological (bright-filed microscopy and CLSM) and physiological analyses (pairing stability, egg production, motility, EdU assays). Finally, we performed RT-qPCR analyses of selected marker genes to substantiate the observed phenotypic effects.

### Maintenance of the *Schistosoma mansoni* life cycle

The *S. mansoni* life cycle was maintained in a controlled environment using *Biomphalaria glabrata* snails as intermediate hosts and Syrian hamsters (*Mesocricetus auratus*) as definitive hosts. Snails were either infected with a single miracidium to obtain unisexual (single-sex; ss) populations of cercariae, or with 10-15 miracidia for mixed-sex (bisex; bs) populations of cercariae. At day 46 (for bs) or 67 (for ss) after infection with the paddling method [102–103], hepatoportal perfusion of hamsters was performed to obtain adult worms and eggs. Worms were cultured *in vitro* in 3 mL M199 medium (Sigma-Aldrich) supplemented with 1% HEPES buffer (1 M), and 1% ABAM-solution (antibiotic-antimycotic) and 10% newborn calf serum at 37°C with 5% CO_2_, as described before [42, 44, 60].

### Ethical standard

Experiments using Syrian hamsters (*Mesocricetus auratus*) as hosts were carried out in accordance with the European Convention for the Protection of Vertebrate Animals for Scientific and Experimental Purposes (ETS No. 123; revised Appendix A) following the 3R principles, and they were approved by the Regional Council in Giessen, Germany (V54-19c 20/15c no. V7/2023).

### MALAR-Y2H

For the identification of potential interaction partners of Sm*gpcr9*, we made use of the Membrane-Anchored Ligand and Receptor Yeast Two-Hybrid system (MALAR-Y2H) [57]. This system allows the detection of protein-protein interactions involving transmembrane proteins [104]. Prey plasmids were transformed into the Y187 strain, and bait plasmids were transformed into the AH109 strain, as described elsewhere in detail [57]. In short, transformed yeasts were selected by plating on synthetic dropout medium (SD) lacking tryptophane (SD/Trp^−^) or leucine (SD/Leu^−^). Mating was performed by resuspending 10 μL of each AH109 and Y187 clone in 500 μL YPDA medium and cultivation for 16 h at 30°C. Yeast hybrids carrying both plasmids were selected on SD/Trp^−^ Leu^−^ plates. Growth assays were done by plating a dilution series (OD600 = 1, 0.1, and 0.01) of two mated yeast clones (n = 2) on SD/Trp^−^ Leu^−^ His^−^ Ade^−^ plates. Interaction was documented after 72 h at 30°C. For β-galactosidase assays, colonies of mated cells were cultivated in SD/Trp^−^ Leu^−^ medium until OD600 reached 0.4 - 0.8, then 1 mL medium was centrifuged, and the cell pellet was dissolved in 400 μL Z-buffer (60 mM Na_2_HPO_4_, 40 mM NaH_2_PO_4_, 10 mM KCl, and 1 mM MgSO_4_, pH 7.0). Afterwards, the cells were lysed by three freeze/thaw cycles in liquid nitrogen. The lysate was dissolved in 200 μL buffer containing 0.4% o-nitrophenyl-β-D-galactopyranoside (ONPG), followed by incubation for 30 min at 30°C. After centrifugation, we measured the absorbance of the supernatant at 405 nm and calculated the Miller units according to the equation: Miller Units = 1000 × OD405/t (min) × OD600.

### *In vitro*-culture conditions and RNAi

Worm couples were taken into culture at the day of perfusion and immediately transferred to supplemented M199 medium, as described before [32, 44]. RNAi experiments of the genes of interest [Smp_244240 (Sm*gpcr9*)] and neuropeptides [Smp_071050 (Sm*npp26*) and Smp_200800 (Sm*npp41*)] were conducted for a period of two weeks. KD efficiency was analyzed by RT-qPCR, and all experiments were performed in triplicates. For RNAi, a two-probe (per gene) RNAi approach was established to successfully knockdown Sm*gpcr9*. To this end, two dsRNAs of this gene were synthesized from different sites of its coding sequence. The dsRNAs were PCR-amplified using gene-specific primers containing the T7 promotor sequence (CCTAATACGACTCACTATAGGGAGA) (S1 Table). After performing a PCR clean-up (Monarch^®^ PCR & DNA Cleanup Kit, NEB, T1030S), each of the respective PCR products were used as templates for the second round of PCR resulting in amplicons that served as templates for the *in vitro* transcription of Sm*gpcr9.* For RNAi of Sm*npp26* and Sm*npp41*, we performed the classical single dsRNA approach [44, 60, 63]. Each dsRNA was synthesized from 300-550 bp long PCR amplicons. As control, we used *ampR* (*Escherischia coli*, ampicillin resistance gene) dsRNA, which has been shown before to be a suitable control for RNAi experiments with adult *S. mansoni in vitro* [63]. The reaction mixture contained 10 µL 10× reaction buffer (0.4M Tris, pH 8, 0.1M MgCl_2_, 20 mM spermidine, 0.1 M DTT), 5 µL PCR product (5 µg), 20 µL 25 mM rNTP (NEB, N0450S), 3 µL self-made T7 RNA polymerase, 1 µL inorganic pyrophosphatase (IPP) (NEB, M0361). The mixture was filled up to a final volume of 100 µL using diethyl decarbonate (DEPC) water. This reaction mixture was incubated overnight at 37°C followed by DNase I (5 µL) (2 U/µL, NEB, M0303) treatment for 30 min at 37°C. Subsequently, the mixture was precipitated with 7.5 M lithium chloride (LiCl) at -80°C for 1 h. After spinning down for 30 min at maximum speed at 4°C, the pellet was resuspended in 70% ethanol. Finally, the mixture was spinned down for 20 min at maximum speed at 4°C. The pellet was resuspended in an appropriate amount of DEPC water, and dsRNAs were stored at -20°C until further use.

For RNAi experiments *in vitro*, worm couples were treated with 15 µg/mL each of the two dsRNAs of Sm*gpcr9*, and 7.5 µg/mL each of the respective dsRNAs of Sm*npp26* and Sm*npp41*. Treated couples were maintained *in vitro* for two weeks with daily monitoring. Separated worms were discarded, and only paired worms considered for further analysis. Untreated (no dsRNA) worms and *ampR* dsRNA-treated worms were used as controls [63]. Culture medium and dsRNAs were refreshed every second to third day. Phenotypic effects of dsRNA-treated worms in culture were monitored by bright-field microscopy focusing on attachment capacity (ability to attach to the petri dish), motility, pairing stability, and egg production. After the experimental period of 15 d, worms were frozen for RNA isolation or fixed for CLSM analyses. All experiments were performed in triplicates.

### RNA isolation, cDNA synthesis, and RT-qPCR analysis

RNA isolation was carried out using Monarch^®^ Total RNA Miniprep Kit (NEB) and transcribed into cDNA with 100-150 ng of total RNA in a single reaction using QuantiTect Reverse Transcription Kit (Qiagen) following the manufactureŕs instructions. RNA was quantified using spectrophotometer and 2100 Bioanalyzer instrument (Agilent Technologies, California, USA). Transcript levels were determined by RT-qPCR diluting the cDNAs 1:10 in nuclease-free water. Experiments were performed using 2x KAPA SYBR^®^ FAST Universal (Roche, KK4618).

Specific primers were designed to give an amplicon of 130-180 bp with a melting temperature of 55-60°C (Primer3Plus; https://www.primer3plus.com) (S2 Table). Primer efficiencies were determined as described elsewhere [68]. In short, primers were designed spanning exon–exon junctions, and melt curve analysis was performed after each amplification run. In all cases, melt curves showed a single distinct peak, confirming amplification of a single specific product. KD efficiencies were calculated using the 2^-ΔΔCt^ method and showed >90% reduction in target gene expression. Primer efficiency was determined using standard curves generated from serial dilutions of cDNA and was between 90-110% in all cases.

The PCR conditions were optimized as follows: initial denaturation at 95°C for 3 min, followed by amplification of 40 cycles at 95°C for 10 s, 60°C for 15 s, and 72°C for 20 s, each with a final extension at 72°C for 2 min. The total reaction volume was 20 µL, and all reactions were performed in triplicates. The expression values were determined with a modified 2^-^^Ct^ method [105], using Sm*letm1* (Smp_065110) as control for normalization. Sm*letm1* was shown before to be a suitable reference gene for gene expression studies with *S. mansoni in vitro* [68].

### Cloning procedures

The complete coding sequence of Sm*gpcr9* was obtained from the WormBase Parasite database (https://parasite.wormbase.org/Schistosoma_mansoni_prjea36577/Info/Index/;55). To enhance efficient expression in mammalian cells, the sequence was human codon-optimized using the GenScript online tool (Tool Version Beta 1.0). This optimization ensured enhanced translation efficiency in HEK293T cells [106]. The human codon-optimized Sm*gpcr9* had an improved GC content of 52%. To aid in the subsequent protein localization and visualization in transfected cells, the Sm*gpcr9* coding sequence was tagged with the red fluorescent protein (dsRed) at the N-terminus and mCitrine, as part of the modified pcDNA3.1(+) (Invitrogen). This construct was subcloned via *Nhe*I (5’) and *Hind*III (3’) into pcDNA 3.1 with mCitrine at the C-terminus, as described before [106]. The integrity of the plasmid construct was confirmed by Sanger sequencing across the dsRed-SmGPCR9 and SmGPCR9-mCitrine junctions (Microsynth).

### Cell culture, transfection, and agonist-induced receptor internalization experiments

Human embryonic kidney 293 (HEK293-6E) cells, licensed from the National Research Council Canada (NRC file 11565, HEK 293EBNA1-6E cell line), were cultured in FreeStyle™ 293 medium supplemented with Pluronic F-68 (colliphor 188) and maintained for at least two passages prior to experimentation [107]. Cells were incubated at 37°C in a humidified atmosphere containing 5% CO₂. Prior transfection, cell viability was assessed using the TC20™ Automated Cell Counter (Bio-Rad), with a viability threshold of >97% considered sufficient for transfection.

Transient transfections were performed using polyethyleneimine (PEI) with a DNA:PEI ratio of 1:3. DNA was diluted in culture medium at a final concentration of 1 µg/mL. For each transfection, DNA and PEI solutions were prepared separately. For a total transfection volume of 2 mL, 6 µg of PEI was diluted in 100 µL of culture medium in one microcentrifuge tube, while 2 µg of plasmid DNA (pcDNA_mCitrine_GPCR9_dsRed) was diluted in 100 µL of culture medium in a separate tube. Each solution was vortexed briefly three times (2 s each), after which the PEI and DNA solutions were combined and gently mixed. The resulting mixture was incubated at room temperature for 5 min prior to addition to the cells. Transfected cells were incubated overnight at 37°C under shaking conditions. Image acquisition was performed using fluorescence microscope (Leica DM IL LED Fluo).

Forty-eight hours post-transfection, cells were treated with 10 µM of neuropeptide agonists (Biomatik, Canada) for 1 h at 37°C. As a positive control, the muscarinic acetylcholine receptor was transfected and stimulated with 100 µM carbachol. Cells were fixed, mounted onto coverslips coated with poly-L-lysine, and visualized using CLSM (TCS SP5 vis confocal laser scanning microscope, Leica) to assess receptor internalization and subcellular localization [108].

### BRET assays and measurements

Experiments were performed using HEK293T cells to investigate Gαq recruitment to the Sm*gpcr9* receptor. To this end, the cells were maintained in Dulbecco’s Modified Eagle Medium (DMEM; Gibco, Thermo Fisher Scientific) supplemented with 10% fetal bovine serum (FBS; Gibco) and 1% penicillin-streptomycin (Pen-Strep; Gibco) at 37°C in a humidified incubator with 5% CO₂.

Transient transfection was carried out using polyethyleneimine (PEI; linear, 25 kDa, Polysciences Inc.) at a DNA (pcDNA_mCitrine_GPCR9_dsRed): PEI weight ratio of 1:3. Prior to transfection, cells were detached using 0.05% trypsin-EDTA (Thermo Fisher Scientific), with a maximum trypsinization time of 3 min. Trypsinization was terminated by adding 5 mL of complete medium per plate. Afterwards, the cells were collected by centrifugation at 1,000 rpm for 3 min at room temperature (RT). The supernatant was discarded, and the cell pellet was suspended in 3 mL of fresh medium per plate.

Cell density was determined using a hemocytometer (Neubauer chamber, depth 0.1 mm). Approximately 14 µL of the cell suspension was used for counting. For transfection, 100 µL of the mixture (containing the DNA:PEI, cells, and medium) was dispensed into each well of a 96-well white-walled plate. Plates were incubated for 48 h at 37°C with 5% CO₂.

Afterwards, cells were washed twice with FRET buffer (e.g., HBSS supplemented with 20 mM HEPES, pH 7.4). Coelenterazine H (Nanolight Technology) was used as the substrate for *Renilla luciferase* II (RlucII) [109–111]. Cells were incubated with coelenterazine H in the dark (being light sensitive) at RT for 10 min.

BRET (Bioluminescence Resonance Energy Transfer) measurements were performed using a TECAN Spark 20M plate reader (SparkControl software) equipped with dual-emission detection, with filters set at 410 ± 80 nm for RlucII (donor) and 515 ± 30 nm for rGFP (acceptor). After acquiring five basal BRET cycles, cells were stimulated with varying concentrations of neuropeptide ligands (NPPs) ranging from 1 µM to 0.01 nM. Emission signals were monitored for 35 cycles, approximately 1 h in total. Each ligand concentration was tested in triplicate. Ligand-induced BRET ratios were calculated by normalizing the emission intensity (rGFP/RlucII) during stimulation cycles relative to the basal signal.

### Molecular docking

Molecular docking studies were performed using the Molecular Operating Environment (MOE 2024.06) software package (Chemical Computing Group, Montreal, CA). SmNPP26b and SmNPP41 were modeled using the MOE Conformational Search in lowModeMD with the following settings: rejection limit 100, iteration limit 100, RMS gradient 0.005, MM iteration limit 500, MM iteration limit 500, RMSD limit 0.25, energy window 7, and conformation limit 200. SmNPP26b generated 1 conformer, and SmNPP41 generated 9 conformers, and these conformers were used as the ligands for docking studies. A homology model of SmGPCR9 (Smp_244240) was created based on the human NPP FF receptor 2, PDB: 9M54 [112]. The SmGPCR9 amino acid sequence was aligned to the NPP FF receptor 2 using the BLOSUM64 matrix, revealing 32% similarity and 23% identity. The template structure was corrected for missing atoms or fractional occupancy based on the amino acid sequence, and the final protein was protonated at T = 310 K, pH = 7.3, [NaCl] = 200 mM, using GB/VI electrostatics. This produced ten intermediate models which were scored based on the electrostatic solvation energy, and the structures were energy minimized using the AMBER12:ETH force field. The final model was determined based on the most favorable electrostatic solvation energy. For docking, the extracellular domain of SmGPCR9 was used as the ligand binding site. Initial placement was calculated for 30 poses per molecule using triangle matching with London dG scoring. The top hits were refined, calculating five poses per molecule with flexible drug and flexible receptor (induced-fit modelling) and Affinity dG scoring, which provides an estimated binding free energy in kcal/mol. The final poses were overlayed via superposition to explore conservation and distances measured between residues of SmNPP26b and SmNPP41 docked to SmGPCR9.

### Whole mount *in situ* hybridization (WISH) and imaging

A modified WISH protocol was used to visualize GPCR transcripts [61]. As samples for analyses, *S. mansoni* couples were first separated using 0.25% tricaine (ethyl 3-aminobenzoate methane sulfonate, Sigma-Aldrich) and killed using 0.6 M MgCl_2_ for 1 min. Then they were fixed in 4% formaldehyde in PBSTx for 4 h followed by rinsing twice with PBSTx and stored in 100% methanol at -20⁰C until further use. Worms were rehydrated by incubation in 50% methanol dissolved in PBSTx followed by bleaching for 1.5 h under light. Samples were then rinsed with PBSTx before proteinase K (20 mg/mL, Ambion, AM2546) treatment. WISH was performed with pairing-experienced bisex males (bM), pairing-experienced bisex females (bF), pairing-inexperienced single-sex females (sF), and pairing-inexperienced single-sex males (sM) to localize transcripts of Sm*gpcr9*, Sm*npp26*, and Sm*npp41*. As positive controls, we used Sm*tsp-2* (Smp_335630) [32, 62] and Sm*myst4* (Smp_165360) [32, 42, 44]. Samples were treated with 4% formaldehyde for 30 min. Riboprobes were generated using DTG (digoxigenin)-11-UTP (Jena Bioscience, NU-821-DIGX, Germany). Riboprobe templates were synthesized from gene-specific inserts cloned into the pJC53.2 plasmid (a kind gift of Jim Collins, Texas) using Q5 High-Fidelity DNA Polymerase (40 U/µL, NEB, M0491S). The purified PCR products were used as templates to finally generate the riboprobes by *in vitro* transcription using T3 (Roche, 11031163001) or SP6 RNA polymerases (Roche, 11487671001). The primers used in this study were listed in S3 table. The reaction mixture contained 100-500 ng PCR product, 2 μL 10x transcription buffer (Roche, 11465384001), 1 µL T3 or SP6 RNA polymerase, 2 µL DIG-NTP mix (10 mM ATP, CTP, GTP and 7 mM UTP, 3.5 mM DTG-11-UTP), 0.6 µL Murine RNase inhibitor (40 U/µL, NEB, M0314S), and nuclease-free water (NEB, T2006-1) to make up the final volume to 20 µL. The reaction mixture was then incubated at 28°C for 16 h. Subsequently, 1 µL of RNase-free DNase I (2 U/µL, NEB, M0303S) was added, and the mix was incubated for 20 min at 37°C. An anti-DIG-AP (1:2,000, Millipore Sigma, 11093274910) antibody was incubated in colorimetric blocking solution [7.5% heat-inactivated horse serum (Sigma-Aldrich, #H1138) in TNT] overnight at 4°C for the colorimetric detection and developed with nitro-blue tetrazolium (Roche, 14799526) and 5-bromo-4-chloro-3′-indolyphosphate (Roche, 13513022). Samples were mounted in 80% glycerol under coverslips and imaged.

### Statistical analysis

Statistical analyses of the data were performed using GraphPad Prism 8.0.2 (GraphPad software). Significant differences were determined by t-test and one way ANOVA. Values were indicated as ns, no significance; *p < 0.05, **p < 0.01, and ***p < 0.001.

## Acknowledgements

We thank Natalia Delis and Sandra Engel for their excellent technical assistance. We thank Christina Scheld for maintaining the *Schistosoma* lifecycle. This work was supported by a grant of the Deutsche Forschungsgemeinschaft 8 (DFG, German Research Foundation): GR1549/7-4 (CGG). The striking image was created with bioRender (https://BioRender.com).

## Supporting information

**Fig. S1. Sm*gpcr9* transcript profile in adult *S. mansoni* according to bulk RNAseq, RT-PCR, and scRNAseq analyses. A-B,** Former data from bulk RNAseq analyses of adult *S. mansoni* and their gonads identified two Smp numbers, Smp_080820.1 and Smp_132220.1, for Sm*gpcr9*, based on version 5 of the genome (42). Recent genome updates, versions 7 and 10, provided the new number Smp_244240 for this gene (55; https://parasite.wormbase.org/Schistosoma_mansoni_prjea36577/Info/Index/); **C,** RT-PCR analysis confirmed the testes-preferential and pairing-influenced transcript profile of Sm*gpcr9* in males. **D,** Single-cell RNAseq showed preferential expression of Smp_244240 in testes (Te) and the neuronal cell cluster 3 (Ncc3) of *S. mansoni* males (56). Abbreviations: bM, males with pairing experience; sM, males without pairing experience; bT, testes of bM; sT, testes of sM; bF, females with pairing experience; sF, females without pairing experience; bO, ovaries of bF; sO, ovaries of sF; Ncc, neuronal cell cluster; Te, testes.

**Fig. S2. Y2H interaction studies provided first evidence for SmNPPs as potential interaction partners of SmGPCR9. A,** MALAR-Y2H assay to detect protein-protein interactions between Sm*gpcr9* and neuropeptides (NPPs) of *S. mansoni*. Shown are results of cell growth assays of yeast strain AH109 transfected with plasmids expressing ligand fusion proteins (NPPs), which was mated with yeast strain Y187 transfected with plasmids expressing Sm*gpcr9* (57). Three different OD600 concentrations of diploid yeast cells were dropped onto SD/Trp⁻Leu⁻His⁻Ade⁻ and Trp⁻Leu⁻ plates, which served as growth control. Colony growth was monitored after 48 and 72 h, respectively. **B**, ONPG-assays to determine β-Gal activity of diploid cells (as in A; 57) showed strongest interactions for SmNPPs 2a, 5b, 15b, 26a, 32.2, and 41, whereas 1a and 24 appeared to be weak putative interaction partners. Shown are the mean values of two clones (n = 2; 57).

**Fig. S3. The Sm*gpcr9* coding sequence adapted to human codon usage to optimise HEK293 cell expression.** The Sm*gpcr9* coding sequence (CDS) of *S. mansoni* was adapted to human codon usage to optimise expression in HEK293-6E cells. Shown here are the optimised Sm*gpcr9* CDS (**A**) and the corresponding amino acid sequence (**B**).

**Fig. S4. Agonist-induced HEK293 transfection experiments showed Sm*gpcr9* internalization with selected NPPs.** HEK293-6E cells transiently expressing pCDNA_mCitrine_GPCR9_dsRed were stimulated with the selected, soluble NPPs for 30 min at 37⁰C. Colocalizing signals were observed as internalized dots. SmNPP11a and SmNPP24 failed to show internalization signals. Scale bars = 30 µm. Experiments were performed in two independent biological replicates.

**Fig. S5. Gαq activation by SmGPCR9 following interaction with selected SmNPPs.** Schematic representations of the same Gαq activation assays showing specific protein–protein interactions of SmGPCR9 with SmNPPs (as indicated). In **A**, the tagged version of SmGPCR9 was used and in **B** the untagged version. SmNPPs 26b and 41, which showed highest evidence for interaction, are part of figure 1.

**Fig. S6. Sm*npp26* and Sm*npp41* transcript occurrence in adults and specific neuronal cell clusters** According to the previous bulk RNA-seq study of adult *S. mansoni* and their gonads, Sm*npp26* (Smp_071050; **A**) and Sm*npp41* (Smp_200800; **C**) are mainly transcribed in adult worms but not their gonads, and with a pairing-dependent expression profile in females, with a bias for unpaired females. Single cell RNA-seq data exhibited dominant expression for Sm*npp26* in neuronal cell cluster 4 (Ncc4; **B**), whereas Sm*npp41* dominates in neuronal cell cluster 8 (Ncc8; **D**) (56). Abbreviations: bM, males with pairing experience; sM, males without pairing experience; bT, testes of bM; Ncc, neuronal cell cluster; sT, testes of sM; bF, females with pairing experience; sF, females without pairing experience; bO, ovaries of bF; sO, ovaries of sF.

**Fig. S7. RNA-seq and RT-qPCR-based transcript profiles of Sm*gpcr9*, Sm*npp26*, and Sm*npp41* in adults and gonads.** Shown are transcript profiles of Sm*npp26* **(A)**, and Sm*npp41* **(B)** obtained by bulk RNA-seq analysis of female and male *S. mansoni* and their gonads (42). RT-qPCR confirmed the transcript patterns of Sm*npp26* **(C)**, and Sm*npp41* **(D)** including their pairing-influenced transcriptions in females. Relative expression levels of transcripts were analyzed by the 2^−△Ct^ method (105), using a previously validated reference gene for analysis and the formula: relative expression = 2^−ΔCt^ × *f*, with *f =* 1,000 as an arbitrary factor as described before (68). Abbreviations: bM, bisex males (pairing-experienced); sM, single-sex males (pairing-unexperienced); bT, testes of bM; sT, testes of sM; bF, bisex females (pairing-experienced); sF, single-sex females (pairing-unexperienced); bO, ovaries from bF; sO, ovaries from sF. Average expression (Avg Expr) was based on RPKM (Reads Per Kilobase per Million mapped reads) values. Significant differences were determined by t-test and indicated as: ***P < 0.001, **P < 0.01, *P < 0.05, ns, no significance.

**Fig. S8. Results of WISH control experiments. A,** As positive controls for WISH, we used the tegumentally expressed Sm*tsp-2* (62) and the vitellarium-specifically expressed Sm*myst4* (32); all of which showed expected transcript patterns. **B,** As negative controls, we used sense probes of the genes, which showed no signals upon hybridisation. Scale bars = 200 µm.

**Fig. S9. Knocking-down transcript levels was more efficient for Sm*gpcr9* using a two-probes/per-target (tp/pt) RNAi approach. A,** Compared to the controls (untreated, no dsRNA; *ampR*, irrelevant dsRNA), using a single dsRNA resulted in the reduction of the Sm*gpcr9* transcript level of about 50% after 15 days *in vitro* treatment. **B,** Using the two-probes/per-target approach, KD efficiency of Sm*gpcr9* was 90% ± 3%. For Sm*npp26* RNAi **(C)** and Sm*npp41* RNAi **(D)**, using single dsRNAs in each case resulted in KD efficiencies of 94% ± 2%, and 93% ± 2%, respectively. Significant differences were determined by t-test and indicated as: ***P < 0.001, **P < 0.01, *P < 0.05.

**Fig. S10. RNAi against Sm*gpcr9*, Sm*npp26*, and Sm*npp41* showed no effects on pairing stability but reduced motility.** Untreated *S. mansoni* couples (without dsRNA) and couples treated with irrelevant *ampR* dsRNA served as controls. All worms were kept under the same *in vitro-*culture conditions for 15 d. **A,** We observed no effect for pairing stability. **B,** Compared to the controls, motility was significantly reduced for all target genes (as indicated) but at different time points, with Sm*gpcr9* showing the slowest significant effect on motility following RNAi. There were no distinct effects observed with *ampR* dsRNA treatment as shown in figures. Significant differences were determined by t-test and indicated as: ***P < 0.001, **P < 0.01, *P < 0.05.

**Fig. S11. RNAi against Sm*gpcr9*, Sm*npp26*, and Sm*npp41* caused egg deformation.** Starting between days 9-12 after dsRNA treatment, deformed eggs were produced in all RNAi groups, as indicated, except the control groups (*ampR* and untreated). Even after 15 days, worms of both control groups produced eggs of normal size with visible zygotes (red circles) and normal spines (blue squares). In contrast, eggs of worms of all RNAi groups showed various defects like size reduction, missing spines, and/or no zygotes.

**Fig. S12. RNAi affected gonadal stem-cell proliferation in *S. mansoni* for Sm*npp26* and Sm*npp41,* not for Sm*gpcr9*.** Results of the Image J-based quantification of the amount of EdU-stained, proliferating cells in the gonads of male (testes) and female (ovary) treated with dsRNA against **(A)** Sm*gpcr9* **(B)** Sm*npp26*, and **(C)** Sm*npp41* versus controls (untreated and *ampR* dsRNA treated). Proliferating cells were also quantified in vitellarium of females **(D)**.

**Fig. S13. Transcript profiles of two tektin orthologs of *S. mansoni* according to different RNAseq analyses.** Previous bulk RNA-seq data of adult *S. mansoni* and their gonads (42) showed testis-preferential expression in *tektin a1* (**A**; previous gene number: Smp_147440; according to the new annotation: Smp_343970) and *tektin 2* (**B**; Smp_046410). **C-D,** Single cell RNA-seq data exhibited dominant expression for both tektins in neuronal cell clusters 2-6 and 30 (Ncc), flame cells Fc), and testes (Te) (56). Abbreviations: bM, males with pairing experience; sM, males without pairing experience; bT, testes of bM; sT, testes of sM; bF, females with pairing experience; sF, females without pairing experience; bO, ovaries of bF; sO, ovaries of sF; Te, testes; NCC, neuronal cell cluster; Fc, flame cell.

**Movies S1. RNAi against Sm*npp26* and Sm*npp41* caused morphological changes in adult *S. mansoni*.** Short video clips **(A-E)** of the motility phenotypes of *S. mansoni* couples observed after 6 days of RNAi against the target genes, as indicated, and the controls (untreated, no dsRNA; *ampR*, irrelevant dsRNA).

**S1 Table. Primers used for dsRNA synthesis of Sm*gpcr9*, Sm*npp26*, and Sm*npp41***

**S2 Table. Primers used for RT-qPCR**

**S3 Table. Primers used for WISH**

